# An *Mtb*-Human Protein-Protein Interaction Map Reveals that Bacterial LpqN Antagonizes CBL, a Host Ubiquitin Ligase that Regulates the Balance Between Anti-Viral and Anti-Bacterial Responses

**DOI:** 10.1101/202598

**Authors:** Bennett H. Penn, Zoe Netter, Jeffrey R. Johnson, John Von Dollen, Gwendolyn M. Jang, Tasha Johnson, Yamini M. Ohol, Cyrus Maher, Samantha L. Bell, Kristina Geiger, Xiaotang Du, Alex Choi, Trevor Parry, Mayumi Naramura, Chen Chen, Stefanie Jaeger, Michael Shales, Dan A. Portnoy, Ryan Hernandez, Laurent Coscoy, Jeffery S. Cox, Nevan J. Krogan

## Abstract

Although macrophages are armed with potent anti-bacterial functions, *Mycobacterium tuberculosis* (*Mtb*) replicates inside these innate immune cells. Determinants of macrophage-intrinsic bacterial control, and the *Mtb* strategies to overcome them are poorly understood. To further study these processes, we used a systematic affinity tag purification mass spectrometry (AP-MS) approach to identify 187 *Mtb*-human protein-protein interactions (PPIs) involving 34 secreted *Mtb* proteins. This interaction map revealed two new factors involved in *Mtb* pathogenesis - the secreted *Mtb* protein, LpqN, and its binding partner, the human ubiquitin ligase CBL. We discovered that an *lpqN Mtb* mutant is attenuated in macrophages, but growth is restored when CBL is removed. Conversely, *Cbl*^-/-^ macrophages are resistant to viral infection, indicating that CBL regulates cell-intrinsic polarization between anti-bacterial and anti-viral immunity. Collectively, these findings illustrate the utility of this *Mtb*-human PPI map as a resource for developing a deeper understanding of the intricate interactions between *Mtb* and its host.

## INTRODUCTION

*Mycobacterium tuberculosis* (*Mtb*) infection persists as a leading cause of death worldwide, with an estimated 2 billion people chronically infected and 1-2 million deaths annually (Zumla et al., 2015). The only vaccine, the live-attenuated Bacillus Calmette-Guerin strain developed nearly 100 years ago, offers very limited protection against *Mtb* (Mangtani et al., 2014). In addition, current treatment is cumbersome, requiring administration of multiple, potentially toxic antibiotics over a period of months (Gillespie et al., 2014). Thus, there remains a critical need to elucidate the mechanisms by which *Mtb* disrupts the host immune response in order to both optimize vaccine strategies as well as to explore therapies that promote sterilizing host immunity as an adjunct to antibiotics.

*Mtb* infection begins when airborne bacilli are inhaled and phagocytosed by alveolar macrophages (Torrelles and Schlesinger, 2017). This activates pattern-recognition receptors that bind bacterial constituents, leading to expression of proinflammatory cytokines such as interleukin 1 (IL-1) and tumor necrosis factor alpha (TNF-α) that are important for *Mtb* control (Mayer-Barber and Sher, 2015). Surprisingly, *Mtb* infection also activates secretion of Type-I interferons such as interferon beta (IFN-β) (Manzanillo et al., 2012), a hallmark cellular response to viral infection (McNab et al., 2015). IFN-β induction has been linked to *Mtb’s* ability to perforate the phagosome membrane through its Type VII/ESX-1 protein secretion system (Manzanillo et al., 2012; Novikov et al., 2011; Siméone et al., 2015; van der Wel et al., 2007), allowing communication between the bacterium and the host cytosol. This allows recognition of extracellular *Mtb* genomic DNA by cyclic GMP-AMP synthase (cGAS) (A. C. Collins et al., 2015; Wassermann et al., 2015; Watson et al., 2015; Wiens and Ernst, 2016) which activates interferon regulatory factor 3 (IRF3) to initiate IFN-β transcription. *In vivo*, IFN-β signaling counteracts anti-*Mtb* immunity (Manca et al., 2004; Manzanillo et al., 2012; Stanley et al., 2007), in-part by antagonizing the effects of interleukin-1 (IL-1) mediated resistance (Mayer-Barber et al., 2014). Moreover, elicitation of antiviral programs may serve to impede antibacterial responses of the host.

While cytokines such as IL-1, TNF-α, and interferon gamma (IFN-γ) are critical for activating macrophages to control *Mtb* growth (Mayer-Barber and Sher, 2015), the cell intrinsic mechanisms by which macrophages restrict *Mtb* are incompletely understood (Braverman and Stanley, 2017). For example, macrophages encode an array of activities capable of eliminating bacteria, including phagocytosis and subsequent delivery to toxic lysosomes (Alonso et al., 2007; Shiloh et al., 1999). *Mtb* has evolved the remarkable ability to replicate within this hostile environment as it is not only intrinsically resistant to low pH and oxidative damage (Darwin et al., 2003; Vandal et al., 2009), but can also inhibit fusion between phagosomes and lysosomes (Armstrong and Hart, 1971; Clemens, 1996; Rohde et al., 2007). Likewise, the mechanisms by which *Mtb* thwarts macrophage resistance mechanisms are unclear. Many bacterial pathogens inject secreted effector proteins into host cells, often to directly interact with host proteins and disrupt host cell function (Byndloss et al., 2017). The ability of *Mtb* to permeabilize its phagosome via ESX-1 and communicate with the host cell cytoplasm raises the possibility that it may similarly introduce secreted effectors to target host proteins. While elegant genetic studies have implicated dozens of secreted *Mtb* factors in virulence (Sassetti and Rubin, 2003; Y. J. Zhang et al., 2013), whether these factors directly interact with host molecules to influence macrophage physiology is unknown.

Identifying the set of physical interactions between secreted *Mtb* and human proteins that occur *in vivo* will be crucial in understanding *Mtb* pathogenesis and may provide new approaches to combat infection. However, only a handful of *Mtb*-host protein-protein interactions (PPIs) have been characterized. For example, a previous yeast 2-hybrid screen targeting *Mtb* proteins identified an interaction between EsxH and the human ESCRT machinery (Mehra et al., 2013). Additional focused studies on individual *Mtb* factors have also identified interactions with the host proteins VPS33B and TLR2 (Bach et al., 2008; Pathak et al., 2007). However, the contribution of these interactions to *Mtb* virulence remains unclear either because of the known pleiotropic effects the bacterial factors have on the physiology of *Mtb* independent of the host (Siegrist et al., 2009), or because genetic disruption of the factors involved does not diminish *Mtb* virulence (Grundner et al., 2008).

Unbiased approaches for characterizing PPIs using mass spectrometry represent powerful ways to probe complex biological systems in an unbiased fashion (Beltrao et al., 2010; S. R. Collins et al., 2007; Krogan et al., 2006). Previously, using affinity tag purification combined with mass spectrometry (AP-MS), we have systematically identified novel, and unanticipated, host pathways required for virulence of a number of important human viruses (Shah et al., 2015), including human immunodeficiency virus (HIV) (Jäger et al., 2012), hepatitis C virus (HCV) (Ramage et al., 2015), and Kaposi’s sarcoma herpes virus (KSHV) (Davis et al., 2015). Likewise, by focusing on a set of *Chlamydia trachomatis* virulence factors secreted into host cells, we uncovered a new pathway by which this difficult and persistent bacterial pathogen manipulates host cells (Elwell et al., 2017; Mirrashidi et al., 2015).

Here we report the results of a similar AP-MS study to identify the PPIs between *Mtb* secreted proteins and host factors. Using a combination of MS and bioinformatic analysis, we identified highly-specific physical interactions with host proteins and coupled this with genetics to begin to probe this PPI network. We have not only uncovered a novel secreted *Mtb* virulence factor essential for *Mtb* growth *in vivo*, LpqN, but have also identified the interacting host CBL ubiquitin ligase as a new restriction factor that regulates cell-intrinsic *Mtb* control. Surprisingly, while CBL-deficient macrophages are more permissive for *Mtb* growth, they are <u>resistant</u> to viral replication, indicating a role in modulating intrinsic control of anti-viral versus anti-bacterial responses during *Mtb* infection. Collectively, these findings demonstrate the value of our *Mtb*-human PPI map as a resource to advance our understanding of the complex interaction between *Mtb* and its host.

## RESULTS

### Creation of an *Mtb*-host PPI network

To explore the mechanisms by which *Mtb* disrupts host immune function, we sought to identify physical interactions between secreted *Mtb* factors and host proteins using a systematic proteomic approach. We reasoned that since the *Mtb*-containing phagosome is permeabilized (Manzanillo et al., 2012; Novikov et al., 2011; Siméone et al., 2015; van der Wel et al., 2007), any protein secreted by the bacterium would have the potential opportunity to interact with host proteins in the cell. We began by analyzing highly controlled culture filtrates of *Mtb,* free of detectable cytoplasmic contamination. Using mass spectrometry (MS), we established a high-confidence set of bona-fide secreted proteins (Figure S1A; Table S1). These data were further curated with additional published MS studies of *Mtb* protein localization (Målen et al., 2007). These conservative criteria, requiring direct biochemical evidence of secretion beyond the bacterial cell wall, were employed to minimize inclusion of non-secreted proteins, though it precludes analysis of secreted proteins present below the level of detection by MS. These analyses identified a high-confidence set of 105 secreted *Mtb* proteins, including 69 proteins with canonical Sec-transporter N-terminal signal sequences and 21 ESX-system substrates (Table S1).

We next expressed each of these proteins in human cells as C-terminal Strep-tag fusions and identified co-purifying host proteins by MS. As expected, expression in 293T cells of the Sec-dependent substrates containing their natural signal sequences led to co-localization with the endoplasmic reticulum (Figure S1B). Thus, we expressed the protein predicted to correspond to the signal peptidase-processed form released by *Mtb* during natural infection, and determined that these factors primarily localized to the cytoplasm (Figure S1B). We initially sought to express the entire set of *Mtb* secreted proteins directly in macrophages but, consistent with reports from other groups (X. Zhang et al., 2009), found this to be an inefficient process that yielded insufficient protein for MS analysis (Figure S1C). To circumvent this issue, yet still allow capture of macrophage-specific factors, we developed a two-step “pull-down” strategy (Figure 1A). After initially expressing and isolating the bacterial factors from 293T cells on Strept-actin resin, the immobilized proteins were then incubated in an *in vitro* binding reaction with lysate from differentiated human U937 macrophages before subsequent AP-MS. We found this method to be robust, with the ability to readily express and purify 99 of the 105 bacterial proteins (Figure S1D). We analyzed a subset of bacterial factors both with and without addition of U937 lysate, and found a substantial number of additional interactions specific to the macrophage lysate (Table S2).

**Figure 1.**
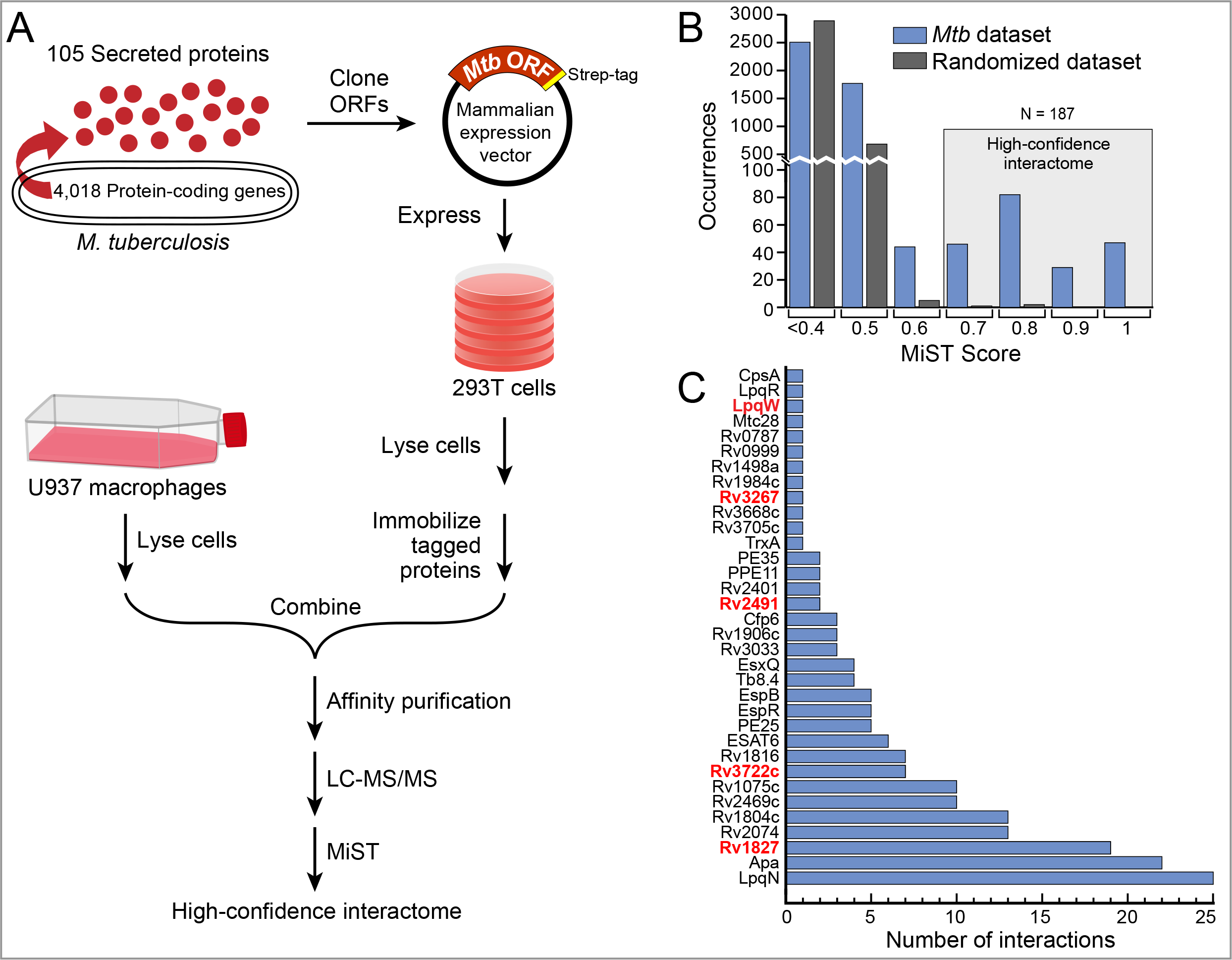
AP-MS experimental design and analysis. (A) Flow-chart of the AP-MS proteomics pipeline used for this study. (B) Distribution of MiST scores for all of the interactions in the *Mtb*-host PPI network, as well as for a randomized control network. (C) Number of high-confidence host interactors (MiST score >0.7, x-axis) for each of the 34 bacterial proteins that had at least one host interaction (y-axis). Genes encoding bacterial proteins likely essential for *Mtb* growth are denoted by red font (Griffin et al., 2011). See also Figure S1.

**Figure 2.**
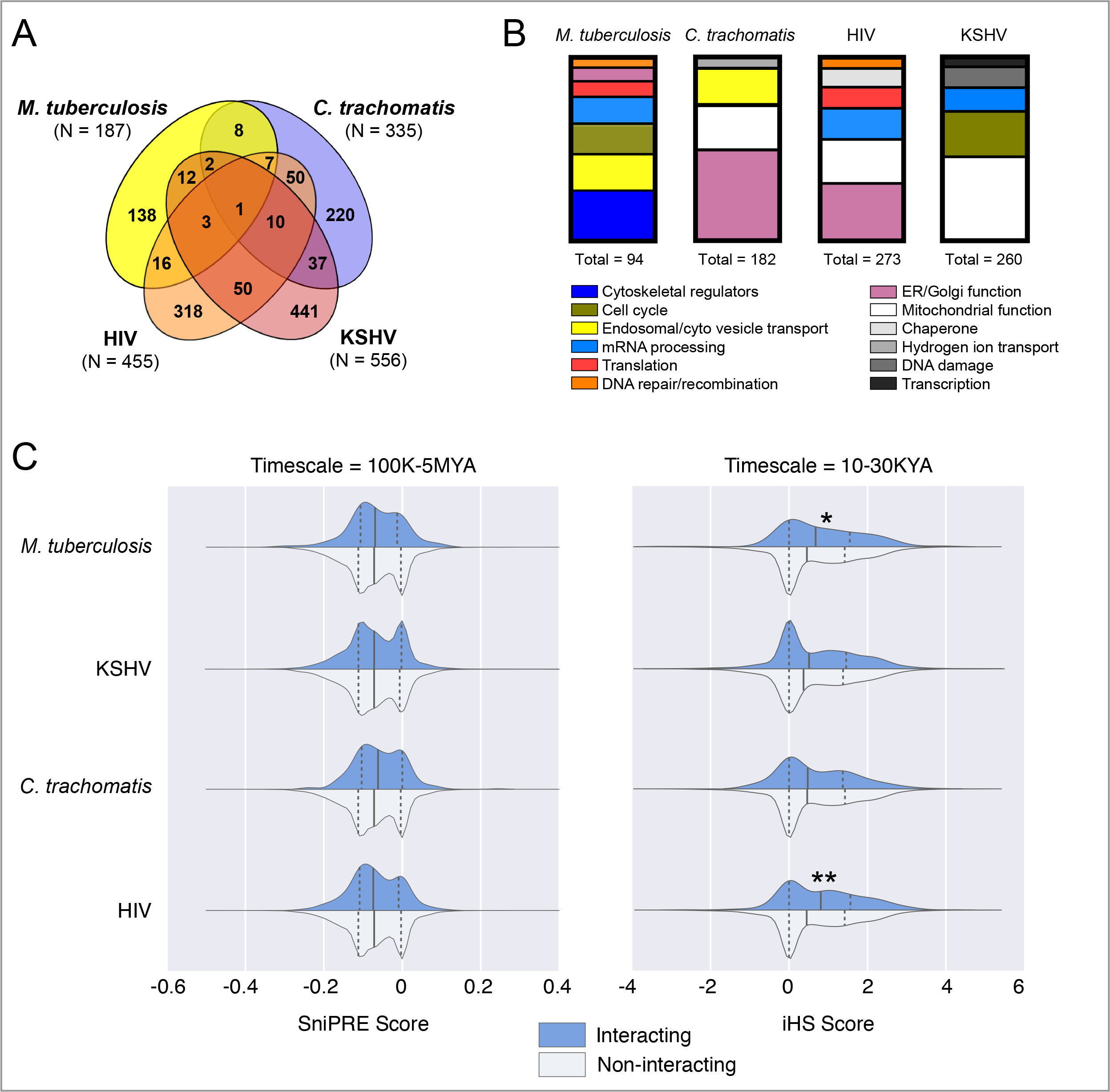
Secreted *Mtb* proteins interact with a distinct, rapidly-evolving set of human proteins. (A) Venn diagram representing the overlap of human proteins identified in four pathogen-host PPI maps. The significance of the overlap in the PPI networks between pairwise comparisons of *Mtb* with HIV, *C. trachomatis*, and KSHV is p=1×10^−9^, p=2.1×10^−5^ and p=0.008 respectively, using Fisher’s exact test. This data is also presented in Table S7. (B) Host biological processes enriched in each PPI map for *Mtb*, *C. trachomatis*, HIV, and KSHV. Processes enriched >2-fold with p-value <0.05 after Benjamini-Hochberg multiple comparisons correction, are displayed. (C) Evolutionary rates of *Mtb*-interacting proteins within the chimpanzee lineage using SniPRE (left) and within the human lineage using iHS analysis (right). The distributions of evolutionary scores across host proteins partitioned into two groups: *Mtb*-interacting proteins shown in blue and non-interacting proteins shown in white, with higher values indicating diversifying selection. Analysis of HIV, *Chlamydia* and KSHV interactomes are shown for comparison. Solid line denotes median, dashed lines denote upper/lower quartile. *p=0.02, **p<0.01.

To identify the set of interacting human proteins, we performed AP-MS in triplicate for each bacterial protein with the addition of GFP-Strep as a control (see Table S3 for raw MS data). To identify specific interactions, we utilized the MiST bioinformatic algorithm, which scores each interaction for specificity, reproducibility, and abundance using principal component analysis (Verschueren et al., 2015). Since no ‘gold-standard’ set of known host-*Mtb* protein interactions existed with which to optimize a score threshold, we used the previously-validated score threshold of 0.7 to define high-confidence interactions (Jäger et al., 2012). Using MiST, the initial interaction dataset was distilled to a high-confidence network between 34 bacterial proteins and 187 specifically-interacting human proteins (Figure 1C; Figure 3; Table S4). As a control, when the AP-MS data were randomized and then analyzed by MiST, only 3 human proteins exceeded the MiST 0.7 threshold (Figure 1B), providing further support that we defined a high quality set of PPIs. To independently validate these interactions, we defined a high-priority set of 34 host interactors with annotated immune-related functions and expressed them in 293T cells as 3×FLAG-tagged proteins with their Strep-tagged bacterial partner. Reciprocal co-immunoprecipitation of the tagged human proteins with their putative bacterial interaction partner verified 25 of 34 (74%) of the interactions, underscoring the reliability of AP-MS coupled with bioinformatic analysis (Figure S1E).

**Figure 3.**
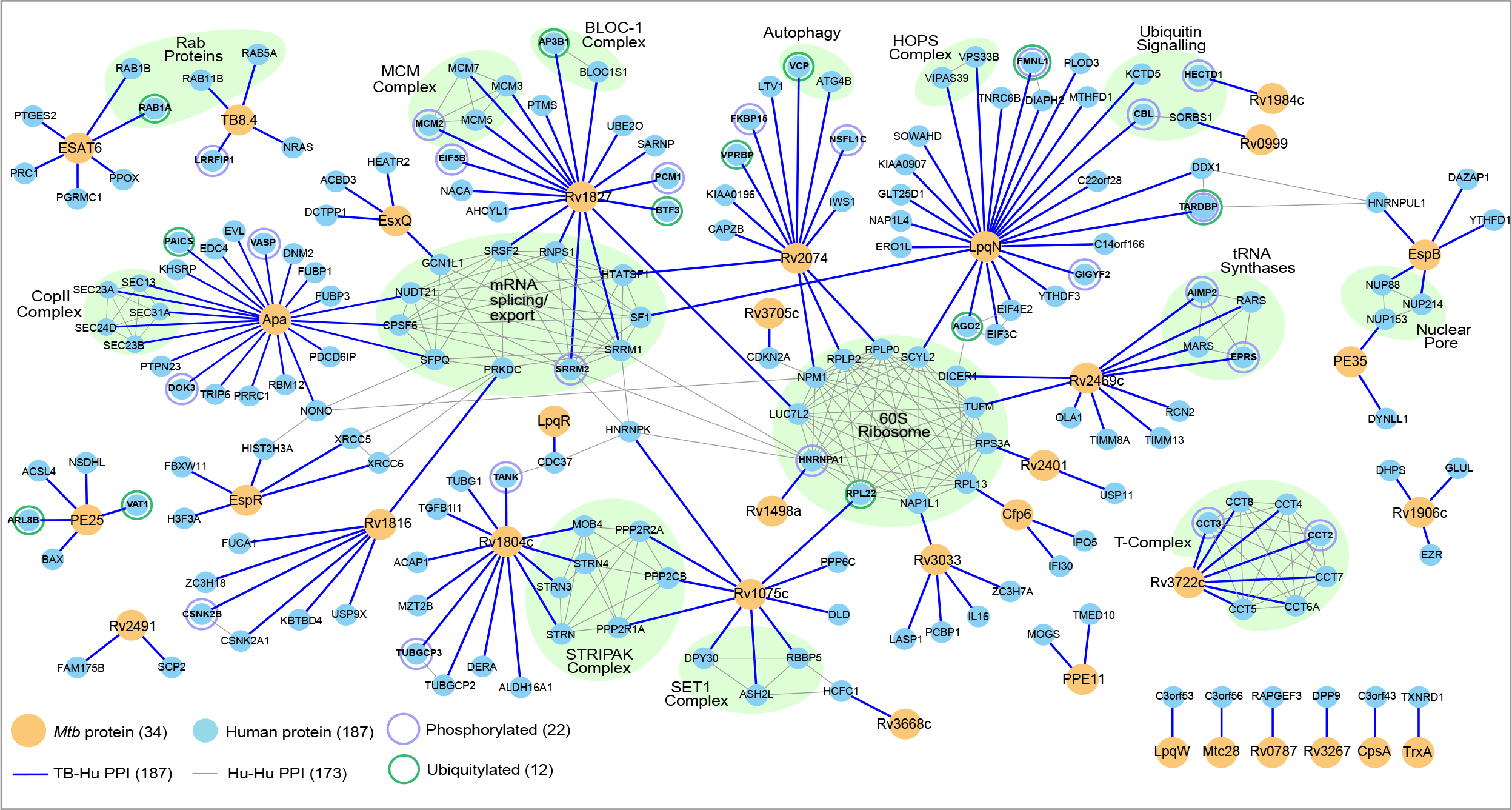
*Mtb*-host protein-protein interaction network. A network representation of the 34 *Mtb* proteins (yellow circles) and 187 human proteins (light blue circles), with blue edges representing the interactions identified in this study. Human-human interactions (thin grey lines) were defined by the CORUM and STRING databases, with known protein complexes highlighted in green. Proteins differentially phosphorylated and/or ubiquitylated upon *Mtb* infection are indicated by concentric circles.

### Features of the *Mtb*-Human PPI Map

Bioinformatic analyses of the interactome revealed several striking features of the *Mtb*-host interaction. We found that only 34% of the *Mtb* proteins we targeted provided high-scoring PPIs with host proteins, which is in contrast to our previous findings with viral proteomes – HIV (89%), HCV (100%), and KSHV (75%) (Davis et al., 2015; Jäger et al., 2012; Ramage et al., 2015). This suggests that, unlike viruses that are completely dependent on host functions for replication, many of the *Mtb*-secreted proteins are involved in bacterial cell-autonomous processes. While we found *C. trachomatis* also had a high percentage of proteins with host interactions (66%), this could reflect the fact that the subset of virulence factors investigated was enriched for mediating contacts with host cells (Mirrashidi et al., 2015). Second, comparison of the host-pathogen interactomes of *Mtb*, HIV, KSHV and *C. trachomatis* revealed that while there was some overlap for the host proteins bound by the different pathogens, the majority of proteins in each of the datasets were pathogen-specific. In particular, 138 of 187 host proteins in the *Mtb*-host interaction map are *Mtb*-specific (Figure 2A, Table S5). However, comparison of the functional pathways represented by these interactions revealed several commonalities between the pathogens (Figure 2B, Table S6). In particular, both *Mtb* and *C. trachomatis* datasets were enriched for proteins involved in vesicular transport, consistent with their common intracellular lifestyles. Moreover, despite some commonalities between the datasets, the interactomes reflected distinct host pathways, suggesting that these pathogens have evolved unique strategies to establish a replicative niche.

### Evolutionary Analysis of the *Mtb*-host interactome

Our data also offer an opportunity to empirically test evolutionary hypotheses regarding *Mtb* and its host. There are numerous examples of host anti-viral restriction factors that are inhibited by direct binding of viral effector proteins. As a consequence, many of these restriction factors experience selective pressure to diversify rapidly (positive selection) and escape the effects of the viral protein - an example of the “Red Queen Hypothesis” of coevolution (Duggal and Emerman, 2012). We hypothesized that if secreted *Mtb* effectors physically interact with host proteins, then these host proteins might show similar evidence of positive selection. We began by using SnIPRE (Eilertson et al., 2012), which compares the relative rate of non-synonymous substitutions between human and chimpanzee orthologs and detects positive selection over a 5-7 million-year span, but identified no evidence of selective pressure over this timeframe (Figure 2C, left panel). However, examination over more recent evolution within the human lineage itself (10,000-30,000-year span), revealed a significant amount of recent diversification (Figure 2C, right panel). In the human population, positively selected alleles quickly sweep to high frequency before recombination separates these variants from the surrounding genome. The integrated Haplotype Score (iHS) measures haplotype block lengths to identify variant alleles undergoing positive selection (Szpiech and Hernandez, 2014; Voight et al., 2006). In contrast to our findings over longer evolutionary timeframes, iHS detected a shift in the distribution of evolutionary rates, with increased diversification for the set of *Mtb*-interacting host proteins (Figure 2C, Table S7). To control for detection bias in our MS methods, we analyzed the set of non-interacting proteins identified in our affinity purifications and found no such shift. We also analyzed the interactomes of other pathogens and found rapid diversification in HIV-interacting host proteins, but not in the interactomes of pathogens associated with lower mortality – KSHV and *C. trachomatis*. Thus, our findings are unlikely to be the result of a bias introduced by either MS detection or bioinformatic filtering. Rather, our data suggest that the set of *Mtb*-interacting host proteins are rapidly diversifying, potentially as a result of their interactions with *Mtb* proteins.

### Topology of the *Mtb*-host interactome

Individual proteins often associate into larger protein complexes to carry out all cellular processes (Alberts, 1998). In an effort to make the *Mtb*-host PPI map more interpretable, we used previously published host PPI data to arrange the identified host proteins into complexes. To this end, we used data from the CORUM and STRING databases to overlay known human PPIs within the *Mtb* interactome (Figure 3). This analysis revealed a number of host complexes targeted by *Mtb* proteins, including a connection between Apa and five subunits of the COP II vesicular trafficking complex as well as an association with the uncharacterized *Mtb* protein, Rv3722c and seven components of the CCT chaperone complex. Furthermore, we uncovered a connection between Rv1804c and Rv1075c and the STRIPAK signal transduction complex, which has been recently linked to innate immunity and autophagy targeting (Liu et al., 2016; Neisch et al., 2017). Interestingly, Rv1075c also interacts with components of the SET1 histone methyltranferase complex, COMPASS, which regulates transcriptional elongation by RNAPII (Krogan et al., 2003). This connection suggests that *Mtb* may regulate host transcriptional regulation by hijacking this complex using one of its secreted proteins.

Recently, our groups carried out a global analysis of changes in the host proteome with respect to post-translational modifications, including phosphorylation and ubiquitination, in the presence of *Mtb* infection (Table S8). Overlaying this dataset with our PPI map revealed that we uncovered 22 and 12 host proteins with altered phosphorylation and ubiquitylation, respectively (Figure 3). These data suggest that these proteins might both be regulated by the host in response to *Mtb*, and targeted by *Mtb* effectors making them high-priority targets for future study.

### LpqN is a novel *Mtb* virulence factor

To begin to identify the functionally relevant interactions between *Mtb* secreted proteins and macrophages, we used a genetic approach to disrupt bacterial and host components of the PPI map and determined the effects during infection. Initially, we selected a set of 10 bacterial factors whose host interactors had known immune-related functions or were post-translationally modified in response to *Mtb* infection, and we generated or obtained previously-created *Mtb* mutants with these genes disrupted. These mutants were evaluated for growth in primary macrophages, and four strains with impaired growth relative to wild-type *Mtb* were carried forward for further analysis (Figure 4A, data not shown). For each mutant, we created two isogenic strains carrying either a wild-type copy of the disrupted gene under the control of its native promoter on an integrated plasmid, or an empty control plasmid. Each plasmid also contained a unique DNA barcode that allowed us to determine the relative abundance of each of the eight strains within pooled infections. We infected mice with this mixture of strains via the aerosol route, recovered bacilli from lung homogenates at multiple time-points, and quantified the relative proportion of each strain by qPCR (Figure 4A). Three of the mutants competed equally with their cognate “complementation” strains, but the *lpqN::*Tn*himar1* mutant (*lpqN* mutant) was rapidly depleted relative to *lpqN::*Tn*himar1::plpqN* (*lpqN* complemented). Importantly, the *lpqN* mutant competed equally with the complemented strain when the pool was grown in culture (Figure S2), and the two strains grew with indistinguishable kinetics in axenic culture (Figure 4B), demonstrating that the *lpqN* mutant was specifically attenuated in the host.

**Figure 4.**
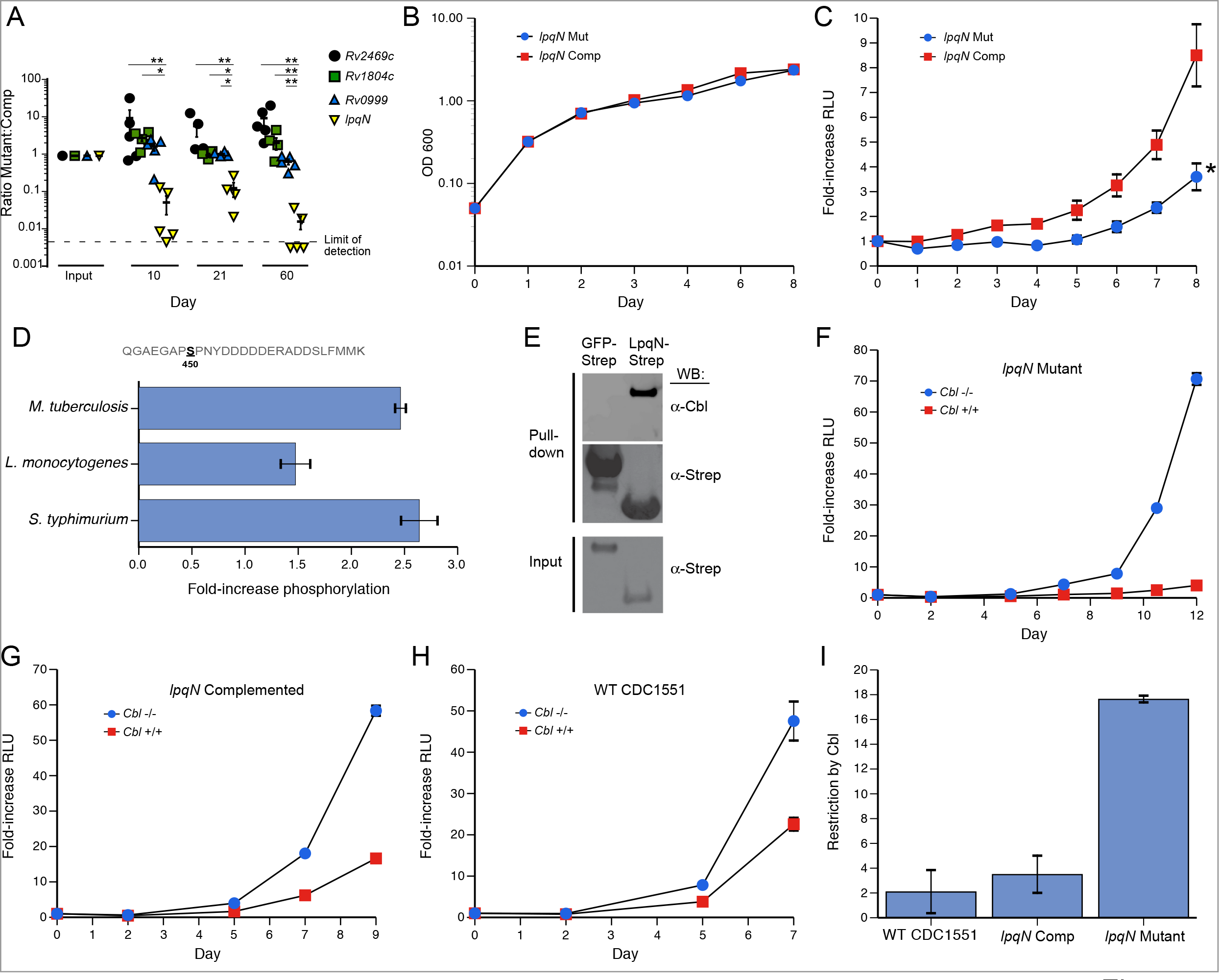
The LpqN-Interacting protein CBL is a host restriction factor for *Mtb*. (A) *In vivo* competition assay. Bacterial mutant strains (LpqN/Rv0583c, Rv2469c, Rv1804c, Rv0999) and cognate complemented strains were pooled and used to infect mice via the respiratory route. At the indicated times, bacteria were recovered from lung homogenates and the relative proportion of each strain was quantified by qPCR using unique sequence tags present in each strain. (B) Growth curve of the indicated strains in standard 7H9 mycobacterial media. Representative data from two independent experiments is shown. (C) Luminescent bacterial growth assay. BMMs were infected with the indicated strains carrying the *LuxBCADE* reporter operon at an effective MOI=1. Relative luminescent units (RLU) were quantified at the indicated times and mean RLU relative to t=0 is plotted. Mean ± SEM are displayed from four replicate samples. Representative data of three independent experiments are shown. * p<0.05 by t-test. (D) Phosphoproteomic analysis. RAW264.7 cells were isotopically labeled and infected with the indicated bacteria. Lysates were analyzed by quantitative LC-MS-MS and the fold-increase for the CBL S450 phosphosite is shown. Mean ± SEM are displayed for two biological replicates, each with two technical replicates. (E) LpqN-Strep or GFP-Strep was expressed in 293T cells and purified with Strep-tactin resin under native conditions, followed by SDS-PAGE and western blotting using antibodies that recognize CBL. (F-H) Luminescent bacterial growth assay. *Cbl*^-/-^ and *Cbl*^+/+^ BMMs were infected with the *lpqN* mutant (F), *lpqN* complemented (G), and WT *Mtb* CDC1551 strain (H). Mean ± SEM of four replicate samples is displayed. Representative data from two independent experiments is shown. (I) Restriction by CBL was derived by determining the ratio of bacterial growth in *Cbl*^-/-^ and *Cbl*^+/+^ BMMs at the final timepoint using data from (F-H). Error bars denote SEM. See also Figures S2, S3 and S4.

We confirmed that the *lpqN* mutant was also attenuated in isolated macrophages by comparing the growth of the isogenic *lpqN* mutant and complemented strains. We used a bioluminescent growth assay in which the strains were transformed with the *luxBCADE* operon from *Vibrio harveyi*, and their growth in macrophages monitored by quantifying luminescence over time. As with our observations in infected mice, we found that the *lpqN* mutant was also significantly attenuated in primary macrophages (Figure 4C), suggesting that LpqN functions during the initial bacterial encounter with the innate immune system. Collectively, these results demonstrate that LpqN is critical for bacterial growth in both *ex vivo* macrophages and in mice, thus establishing LpqN as a novel *Mtb* virulence factor.

### LpqN and the ubiquitin ligase CBL interact physically and genetically

We hypothesized that LpqN functions to promote bacterial growth through its interaction with host proteins, either by blocking their function or by ‘hijacking’ them for pathogenic purposes. To test this idea, we used CRISPR/Cas9-based mutagenesis in RAW 264.7 cells to determine whether disruption of LpqN-interacting host factors was able to rescue the growth defect of the *lpqN* mutant. As noted above, our related studies using MS to examine host protein post-translational modifications during *Mtb* infection demonstrated increased phosphorylation of the ubiquitin ligase CBL, a LpqN-interacting protein, and suggested a possible role for CBL in antibacterial immunity (Figure 4D; Table S8). Indeed, while mutagenesis of several other LpqN interactors failed to rescue the *lpqN* mutant phenotype, we observed that disruption of *Cbl* rescued the growth of the *lpqN* mutant in RAW264.7 cells (Figures S3A and S3B).

We verified the physical interaction between LpqN and CBL by expressing LpqN-Strep in 293T cells. Endogenous CBL protein that copurified was detected by western blotting, verifying the interaction detected by MS (Figure 4E). Additional *in vitro* pull-down experiments using LpqN and CBL proteins produced in *E. coli* also revealed direct interaction between these two factors (Figure S4). We further explored the genetic interaction between *lpqN* and *Cbl* using primary bone marrow-derived macrophages (BMMs) deficient in CBL. To avoid the potential confounding effects of the two related CBL family ubiquitin ligases, we analyzed BMMs lacking the *CblB* and *CblC* genes, and carrying either a wild-type or floxed *Cbl* locus that could be deleted with ~90% efficiency by addition of 4-hydroxy-tamoxifen to the culture media - hereafter designated *Cbl*^+/+^ and *Cbl*^−/−^ respectively (Figure 5I)(Mukhopadhyay et al., 2016). We infected *Cbl*^+/+^ and *Cbl*^-/-^ primary macrophages with the *lpqN* mutant strain and monitored bacterial growth over time. Consistent with our observations in *Cbl*^-/-^ RAW264.7 cells (Figure S3B), the *lpqN* mutant was markedly attenuated in *Cbl*^+/+^ BMMs (Figure 4F). Importantly, in *Cbl*^-/-^ BMMs, we observed significant rescue of the *lpqN* mutant intracellular growth phenotype, with a 15-fold increase in bacterial growth (Figure 4F). This result was further confirmed by plating macrophage lysates and directly enumerating bacterial colony-forming units (CFU) (Figure S3C). We also noted that both the *lpqN* complemented strain and the parental WT CDC1551 *Mtb* strain grew ~2-fold faster in *Cbl*^-/-^ macrophages, indicating that CBL imposes some restriction on wild-type *M. tuberculosis* (Figures 4G, 4H, S3D), however, the effect of CBL activity was much more pronounced with the *lpqN* mutant (Figure 4I). Importantly, the growth of an attenuated ESX-1 mutant of *Mtb*, which is unable to permeabilize its phagosome, was unaffected in *Cbl*^-/-^ macrophages (Figure 5G), as were the growth of other intracellular bacteria including *Listeria monocytogenes* and *Salmonella enterica* serovar Typhimurium (*S.* Typhimurium, Figures S3E and S3F). Thus, there is a genetic interaction between bacterial *lpqN* and host *Cbl*, whereby growth of the *lpqN* mutant strain is selectively enhanced upon loss of host *Cbl*, but strains expressing LpqN are relatively insensitive to CBL activity. This genetic interaction suggests that LpqN and CBL may lie in a common pathway, and collectively the combined biochemical and genetic data suggest a model whereby CBL acts as a host restriction factor, limiting *Mtb* growth, and LpqN acts as a virulence factor to block the normal functions of CBL and promote bacterial replication.

**Figure 5.**
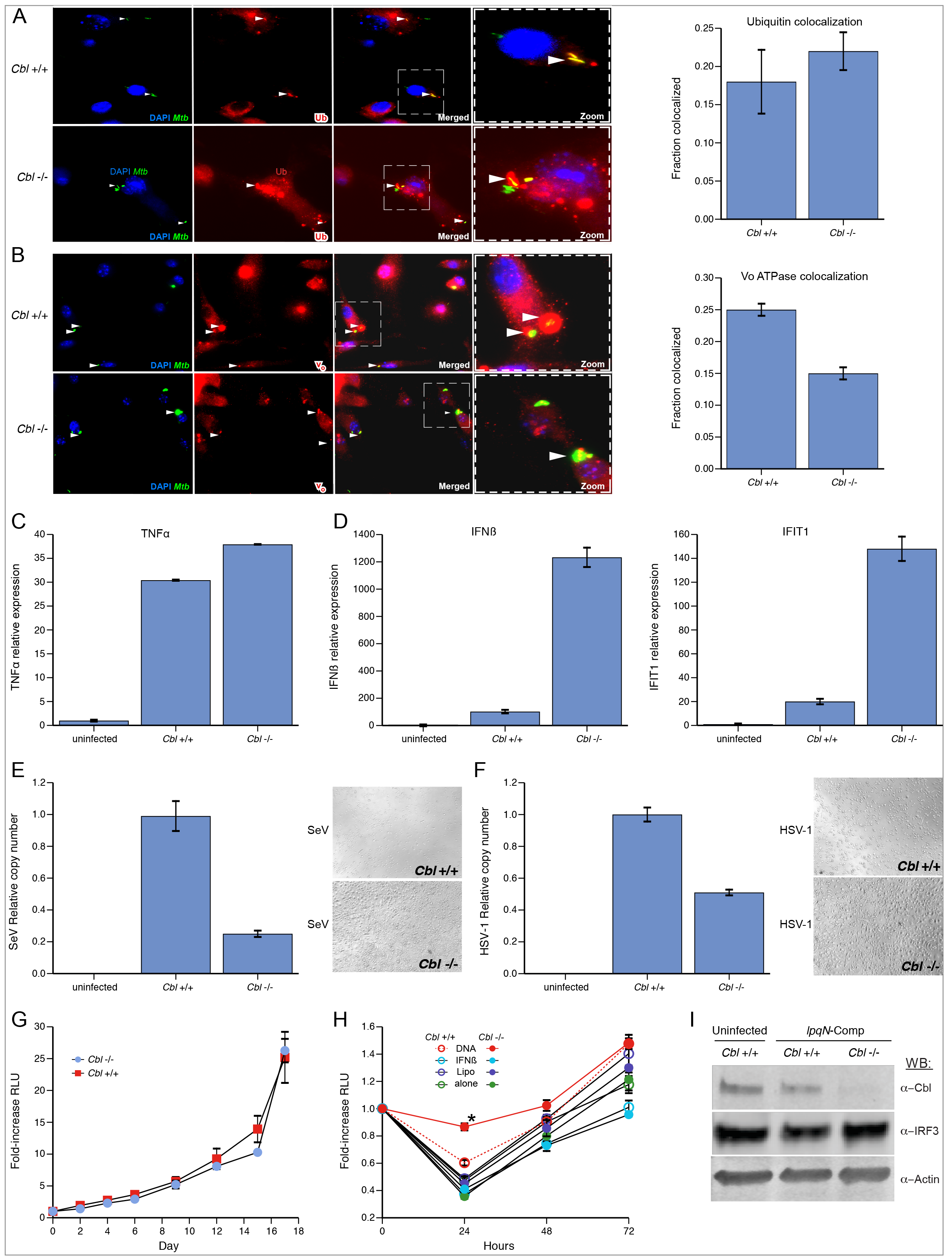
CBL represses anti-viral responses during *Mtb* infection. (A) *Cbl*^-/-^ and *Cbl*^+/+^ BMMs infected with *lpqN* mutant for 4h and immunostained for polyubiquitin. Percent colocaliztion of >500 phagosomes counted per condition, scored by a microscopist blinded to sample identity. Mean ± SEM displayed. (B) BMMs infected with *lpqN* mutant *Mtb* for 24h and immunostained for vacuolar ATPase. (C-D) BMMs infected with *lpqN* mutant *Mtb* for 6 hours and analyzed by RT-qPCR for the proinflammatory cytokine TNF-α (C), or the anti-viral response genes IFN-β and IFIT1 (D). (E) BMMs infected with Sendai virus (SeV) for 24h, and the relative number of virions in the supernatant were quantified by RT-qPCR; Mean ± SEM displayed. Phase-contrast image (10×) taken at 24h post-infection. (F) Infection with HSV-1 for 24h, analyzed as in (E). (G) Luminescent growth assay of ESX-1-deficient strain (*ΔEccC-LuxBCADE*) in BMMs. (H) Luminescent growth assay. BMMs were treated with DNA (Lipo+DNA), transfection reagent alone (Lipo), or IFN-β (250U) as indicated. Mean ± SEM of four replicate samples are shown. IFN-β was added 4h pre-infection and DNA was delivered 1h post-infection. *p=0.003 by t-test (I) BMMs were infected with *lpqN* mutant *Mtb* for 6h and analyzed by Western blot.

**Figure 6.**
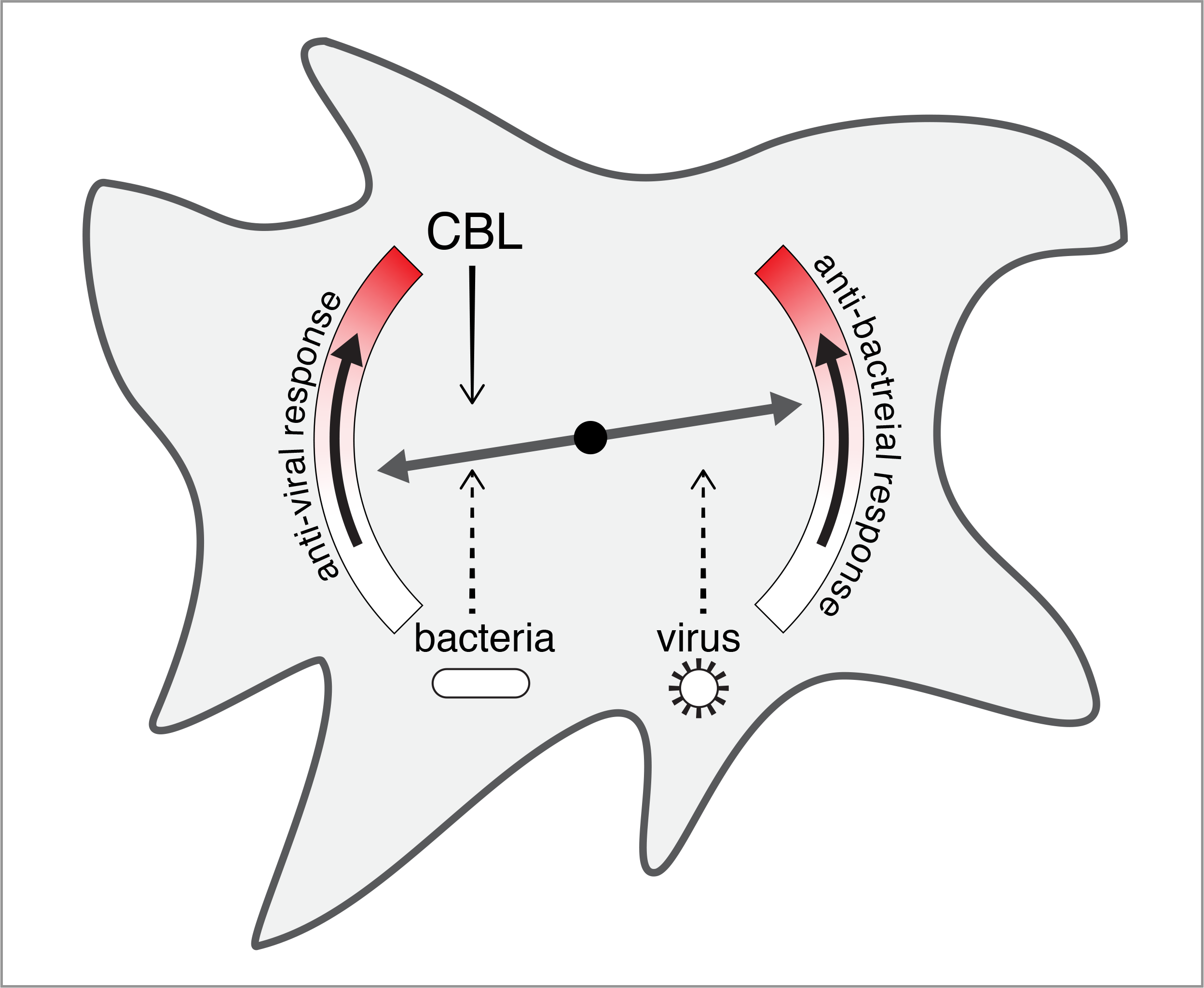
Model of balance between anti-viral and anti-bacterial cell-intrinsic immune response pathways. Host macrophages tailor responses to distinct kinds of pathogens at the earliest stages of infection by activating cell-intrinsic immune pathways tailored to the threat, e.g. virus or bacterium. These programs appear to be mutually antagonistic, as activation of antiviral pathways comes at the cost of antibacterial immunity during *Mtb* infection. Our work indicates that CBL functions to influence this balance by inhibiting viral responses and promoting antibacterial immunity. Macrophage plasticity can be manipulated by pathogens (dashed arrows) to tip the balance to promote inappropriate responses to favor replication.

### CBL regulates the balance between cell-intrinsic anti-bacterial and anti-viral responses

CBL has been well-characterized as a ubiquitin ligase responsible for ubiquitylation and degradation of activated receptor protein tyrosine-kinases (Levkowitz et al., 1999). Since ubiquitylated proteins localize to the *Mtb* phagosome and are proposed to function as a targeting mechanism for autophagy-mediated host defense (Ponpuak et al., 2010; Watson et al., 2012), we initially hypothesized that CBL might directly contribute to this process. The frequency of ubiquitin colocalization with the *Mtb* phagosome after 4h of infection was unaffected in *Cbl*^-/-^ BMMs (Figure 5A). However, to more broadly evaluate the efficiency by which *Mtb* is delivered to the lysosome, we infected *Cbl*^+/+^ and *Cbl*^-/-^ BMMs with the *lpqN* mutant and quantified colocalization of bacteria with the vacuolar ATPase. After 24h of infection, we observed a significant decrease in colocalization with the vacuolar ATPase in *Cbl*^-/-^ BMM (Figure 5B), consistent with the idea that CBL promotes innate bacterial restriction mechanisms of naïve macrophages.

Given the role of CBL as a regulator of signaling, we also investigated whether the ligase modulates the macrophage response to bacterial infection by monitoring expression of key cytokines activated in response to *Mtb*. Though we observed no difference in TNF-α mRNA levels between *Cbf*^+/+^ and *Cbl*^-/-^ cells (Figure 5C), we noted a dramatic increase in the expression of the anti-viral cytokine IFN-β (Figure 5D). Likewise, IFIT1, another target of antiviral signaling, was also increased in *Cbl*^-/-^ cells (Figure 5C). Thus, the ability of *Mtb* to activate antiviral responses, likely through the exposure of bacterial DNA to the cytoplasmic sensor cGAS, is amplified in the absence of CBL, indicating that this ligase functions to negatively regulate these responses.

These data support the notion that CBL-deficient macrophages are skewed towards more robust anti-viral responses, which come at the cost of decreased anti-bacterial resistance. A prediction of this hypothesis is that *Cbl* ^-/-^ cells would be resistant to viral infection. To test this, we infected *Cbl*^-/-^ and *Cbl*^+/+^ BMMs with Sendai Virus (SeV) and Herpes Simplex Virus 1 (HSV-1) and evaluated the release of viral particles 24h after infection. Deletion of *Cbl* conferred resistance to both viruses, with fewer viral particles released, and a reduction in viral cytopathic effects (Figure 5E, 5F). Thus, while loss of CBL results in increased sensitivity to *Mtb* infection, it also creates a more restrictive environment for viral replication. This is consistent with recent findings that siRNA-mediated depletion of CBL results in stabilization of IRF3 and increased anti-viral signaling (Zhao et al., 2016), though when we evaluated whether CBL similarly regulated IRF3 during *Mtb* infection we detected no change in protein levels (Figure 5I). Although the exact mechanism by which CBL influences antibacterial/antiviral balance, the permissiveness of *Cbl*^-/-^ macrophages for *Mtb* growth is unlikely to result from increased Type I interferon itself, as isolated macrophages deficient for Type I IFN production (*Irf3*^-/-^) or reception (*Ifna*r^-/-^) are not altered in their ability to restrict *Mtb* replication (Manzanillo et al., 2012; Stanley et al., 2007). Thus, it appears that CBL may control a broader cell-intrinsic anti-bacterial program by inhibiting anti-viral responses, with one component being Type I IFN activation.

Although the exact role of ESX-1 in *Mtb* virulence remains mysterious, it is generally assumed that the poor growth of ESX-1 mutants in macrophages is due primarily to their inability to perforate phagosomal membranes and access the cytosol to acquire nutrients and deliver effectors (Stanley and Cox, 2013). However, given the reciprocal effects of antiviral and antibacterial responses mediated by CBL, it is also possible that the inability to activate the CBL-regulated antiviral program directly contributes to the attenuated phenotype of ESX-1 mutants. As discussed above, when we infected BMMs with an ESX-1 mutant, we observed no effect of CBL on bacterial growth (Figure 5G). This supports the idea that CBL acts by suppressing the cellular anti-viral response, and that in the absence of an anti-viral signal, CBL is no longer necessary to control the growth of ESX-1 mutants. To further investigate the idea that a CBL-regulated anti-viral program promotes *Mtb* growth, we tested whether ectopic activation of anti-viral responses by experimentally delivering DNA to the host cell cytosol would rescue the growth defects of ESX-1 mutant bacteria. In this way, activation of the anti-viral pathway negatively regulated by CBL would occur in the absence of phagosomal permeabilization, effectively bypassing the ESX-1 requirement. In macrophages transfected with DNA we observed a transient rescue of the ESX-1 mutant with impaired bacterial killing over the first 24h, an effect most notable in *Cbl*^-/-^ macrophages (Figure 5H). This suggests that anti-viral signaling is sufficient to blunt the anti-bacterial activity of macrophages and that CBL functions to counteract this effect. Importantly, addition of IFN-β did not rescue growth of the ESX-1 mutant, consistent with the idea that CBL regulates an anti-viral program that is independent of Type I IFN.

## DISCUSSION

Despite the impact of TB on mankind, surprisingly little is known about the physical interactions between *Mtb* proteins and its human host proteome. Our understanding of how many enteric pathogens interact with host cells via secretion of virulence factors that target host pathways has shed light on the evolution of intricate interactions that underlie the host-pathogen interface. Because *Mtb* is a remarkably successful bacterial pathogen, capable of persisting for the lifetime of the host, it seems likely that *Mtb* would similarly introduce secreted effectors to manipulate host pathways and forestall host immunity. However, the evidence that *Mtb* actually employs this strategy is scant. There are only a handful of characterized interactions between *Mtb* and host proteins, and it remains unclear how they impact *Mtb* virulence. We have addressed this question using our AP-MS pipeline to globally identify 187 high confidence, physical interactions between *Mtb* and host proteins. It is likely that not all of these will contribute significantly to bacterial virulence during the course of a natural *Mtb* infection. However, this proteomic approach, when combined with genetic analysis, represents a powerful way to identify new, biologically relevant interactions between pathogens and their host cells. Indeed, with this approach we have identified a new *Mtb* virulence factor, LpqN, and additionally have identified the CBL ubiquitin ligase as a protein that interacts both physically and genetically with LpqN and functions as a host restriction factor for *Mtb*, limiting bacterial growth in macrophages. We expect that further mining of this interaction map will reveal further interactions that underlie *Mtb* pathogenesis.

CBL had not been implicated in innate responses to bacteria, and the mechanisms by which it contributes to host immunity await discovery. Ubiquitin ligases play key roles in targeting of intracellular pathogens to autophagy, but our data indicates that CBL plays a regulatory role modulating cell-intrinsic responses to infection. Indeed, we noted a surprising increase in the cellular anti-viral response in the absence of CBL, with a concomitant decrease in the efficiency of anti-bacterial processes such as phagosome-lysosome fusion. Importantly, CBL had no effect on the growth of an ESX-1 mutant strain of *Mtb* that does not trigger anti-viral responses. This finding that the anti-bacterial properties of CBL are only manifest in the setting of an anti-viral host response is consistent with the hypothesis that CBL is acting as a regulator of the anti-viral response to control bacterial infection.

The specific antibacterial pathways regulated by CBL in the context of *Mtb* infection remain unknown. IFN-β has been suggested to impair clearance of the related pathogen *Mycobacterium leprae* (Teles et al., 2013), and disruption of either IRF3 or IFNAR have previously been shown to modestly impact *Mtb* growth in mice (Manzanillo et al., 2012; Stanley et al., 2007). However, IFN-β signaling itself does not alter *Mtb* growth cell-intrinsically in isolated macrophages, as we observe with CBL. This suggests additional, uncharacterized antiviral pathways exist, and are activated by *Mtb*, in addition to the known IRF3-IFN-β-IFNAR pathway. The existence of additional anti-viral pathways is further supported by our findings in a related study using proteomics to examine host protein post-translational modifications during *Mtb* infection (Parry et. al. *in preparation*). In that work, we identified the IRF7-regulated antiviral response as a potent facilitator of *Mtb* replication. The effects mediated by IRF7 and CBL share several similarities. Unlike the IRF3-IFNAR pathway, they both alter *Mtb* replication cell-intrinsically in isolated macrophages. The effects of perturbing IRF7 and CBL are also both specific to *Mtb*, as the replication of neither *S*. Typhimurium nor *L. monocytogenes* are altered. It remains to be determined whether CBL and IRF7 are actually functioning in the same pathway, or whether they modulate parallel anti-viral pathways. The finding that two different regulators of the macrophage anti-viral response both impact the ability of *Mtb* to replicate inside macrophages suggests that this is an essential element in TB pathogenesis. We postulate that *Mtb* might have evolved the ability to release bacterial DNA into the host in order to trigger the macrophage anti-viral response, and that this anti-viral response antagonizes antibacterial activity. Effectors such as LpqN could then potentially function to prolong or amplify such a signal and create a more permissive intracellular environment for bacterial growth.

From within its replicative niche in host macrophages, *Mtb* is able to undermine the host immune response and establish a chronic, often lifelong infection – but how it actually does so remains mysterious. The interaction map reported here represents a unique resource to generate testable hypotheses regarding the *Mtb*-host interface and to identify the host pathways that dictate TB pathogenesis.

## AUTHOR CONTRIBUTIONS

B.H.P., N.J.K. and J.S.C. conceived the project; B.H.P., Z.N, J.R.J., T.P, D.A.P., R.H., L.C., N.J.K. and J.S.C. designed experiments; B.H.P., Z.N, J.R.J., J.V.D., G.M.J., T.J., Y.M.O., C.M., S.L.B., K.G., X.D., A.C., T.P., C.C., and S.J. performed the experimental work; B.H.P., Z.N, J.R.J., J.V.D., M.S., D.A.P., R.H., L.C., N.J.K. and J.S.C. analyzed the results; B.H.P., Z.N., C.M., N.J.K. and J.S.C. wrote the manuscript; M.N., D.A.P., R.H. and L.C. contributed reagents and technical expertise.

## ACKNOWLEDGMENTS

We thank members of the Cox, Krogan and Stanley (UCB) labs for comments on the manuscript and for invaluable discussions. This work was supported by NIH grants P01 AI063302 (N.J.K., J.S.C., D.A.P.), P50 GM082250 (N.J.K.), U19 AI106754 (N.J.K.), DP1 AI124619 (J.S.C.), and R01 AI120694 (N.J.K. and J.S.C.). B.P.H. was supported by an A.P. Giannini award and NIH K08 (K08AI104766).

## SUPPLEMENTAL INFORMATION

**Figure S1.**
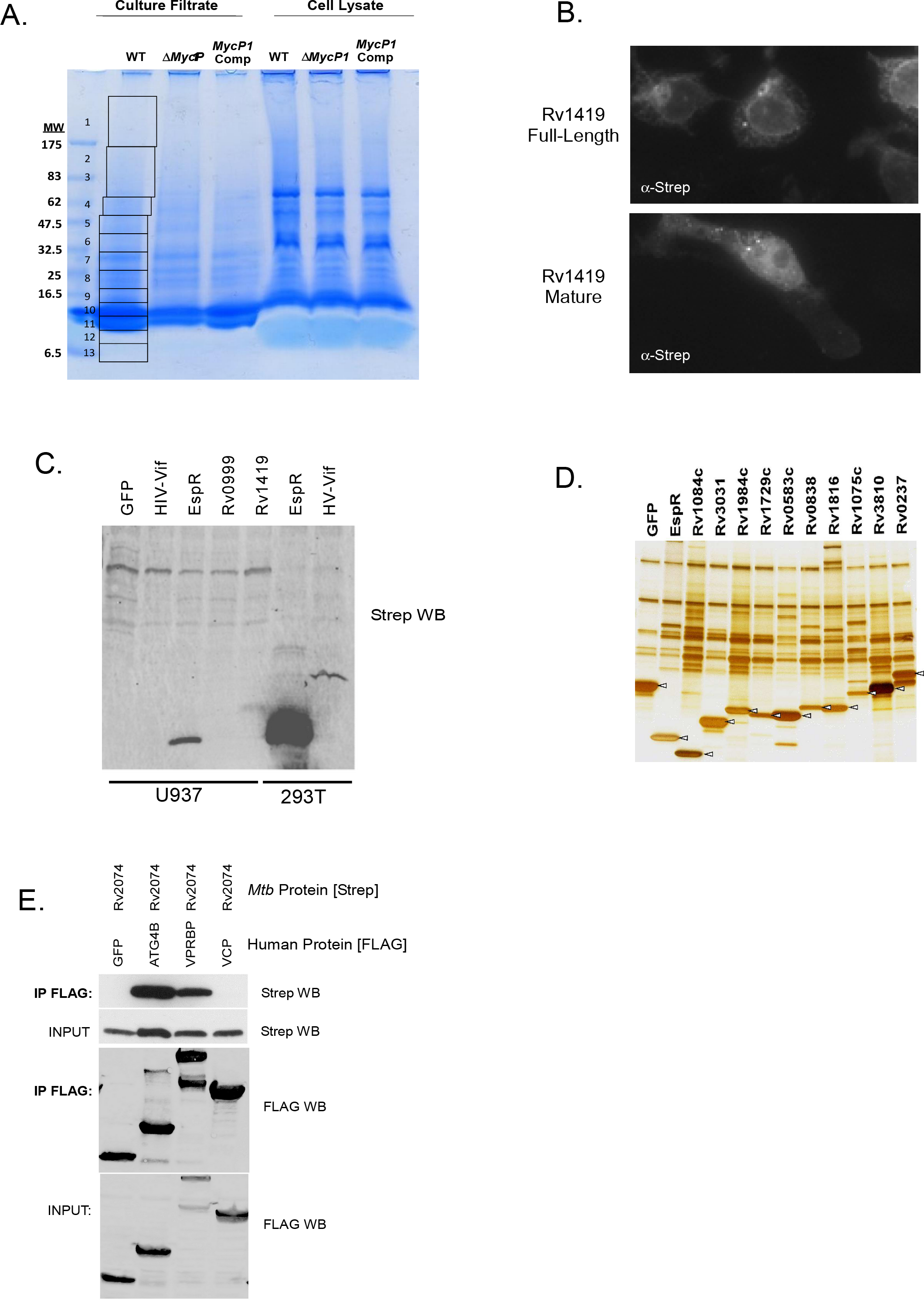
Related to Figure 1. Establishing conditions for proteomic analysis of secreted *Mtb* proteins in human cells. (A) Culture filtrates were prepared from wild-type cultures, separated by SDS-PAGE, and proteins identified by mass spectrometry. The gel was cut into sequential regions as indicated, and each slice was individually subjected to in-gel trypsin digestion and LC-MS/MS (Table S1). Supernatants from ESX-1 mutants (Δ*mycP1*) indicate that most of the secreted proteins are not ESX-1 substrates, and cell lysates were added as controls for lysis. (B) Immunofluorescence microscopy of 293T cells expressing either the full-length uncleaved *Mtb* Rv1419 protein, or the form predicted to be released after signal peptidase cleavage. (C) Western blot analysis of U937 macrophages and 293T cells transfected with plasmids encoding the indicated Strep-tagged proteins, showing much higher levels of expression in 293T cells. The majority of proteins were undetectable from U937 cells, but were expressed to sufficient levels in 293T cells to purify and detect host interactors. (D) Affinity purifications from lysates derived as described in Figure 1 were separated by SDS-PAGE and visualized by silver staining, demonstrating the efficient expression and purification of bacterial proteins. Arrowheads denote the tagged bacterial proteins. (E) Example of reciprocal co-immunoprecipitation validation studies. 293T cells were cotransfected with plasmids expressing Strep-tagged *Mtb* Rv2074 and with FLAG-tagged versions of GFP (negative control), or each of three putative interactors identified in the PPI map, VPRBP, ATG4B, and VCP. The host proteins were immunoprecipitated with anti-FLAG antibody and Rv2074 was detected by Western blot using anti-Strep antibodies.

**Figure S2.**
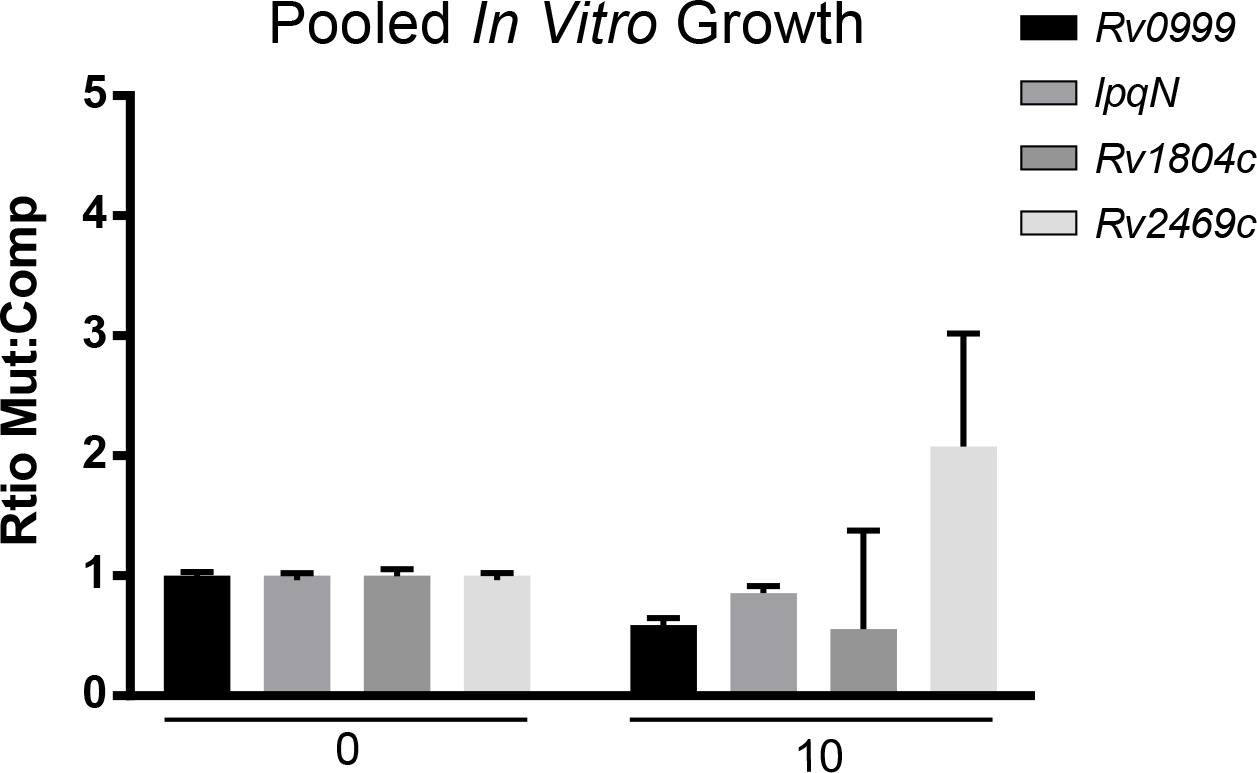
Related to Figure 4. The LpqN mutant competes normally in mixed cultures grown in liquid media. An aliquot of the pool of eight strains used to infect mice for the *in vivo* competition assay (Figure 4) was grown in liquid 7H9 media for 10 days and the relative proportion of each strain was quantified by qPCR of the genomic sequence tag. Mean ± SEM is displayed.

**Figure S3.**
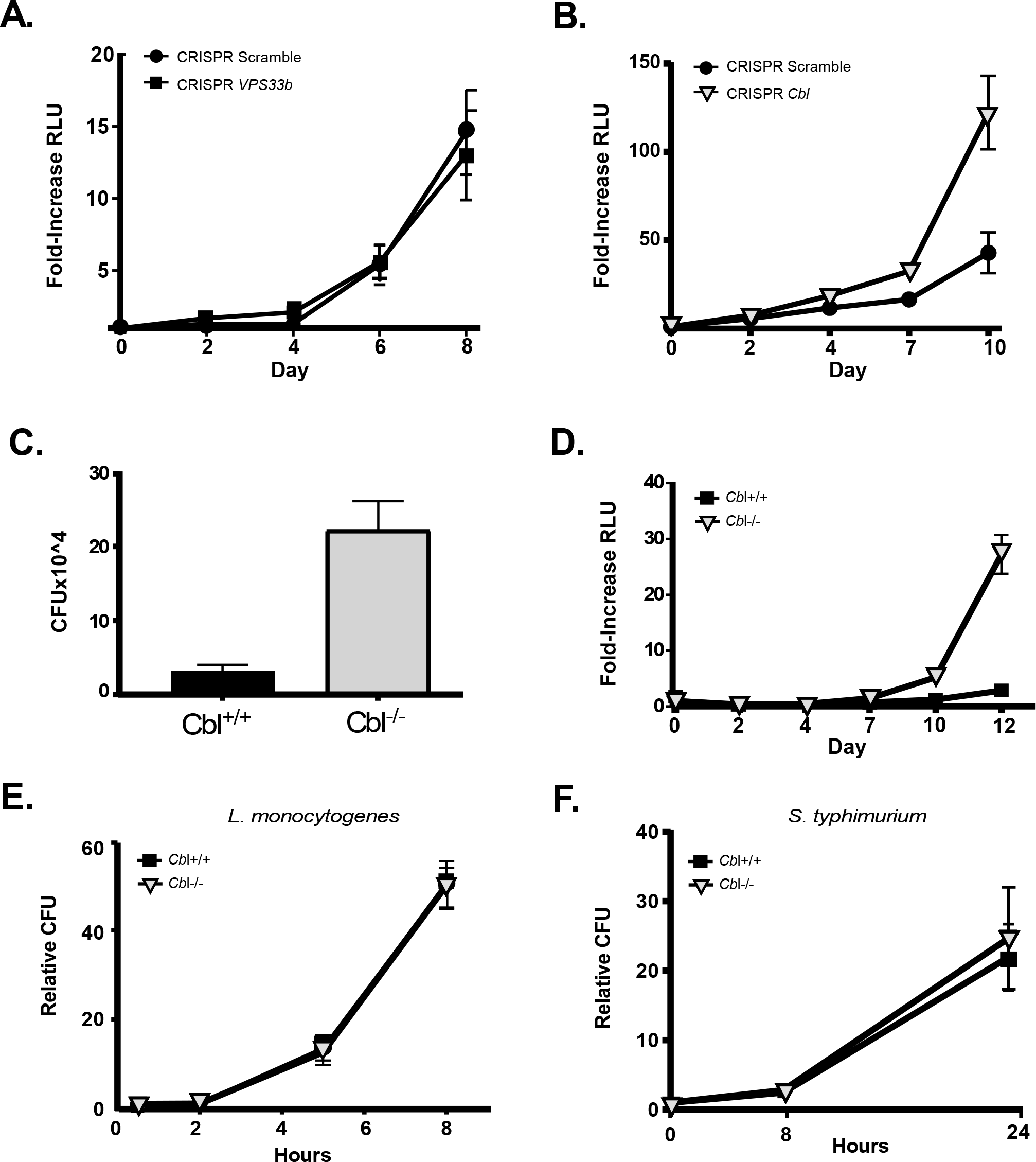
Related to Figure 4. Genetic interactions between Host *Cbl* and *Mtb lpqN* in RAW264.7 cells. (A) The gene encoding VPS33b, a putative LpqN interactor, was mutated by CRISPR/Cas9 in RAW264.7 cells. These cells were infected with the *Mtb lpqN* mutant strain expressing the *luxBCADE* operon from *Vibrio harveyi,* and bacterial growth was assessed by measuring luminescence. The assay was performed twice with three independent mutant clones and two independent wild-type control clones. Each condition was measured in quadruplicate; mean ± SEM from representative clones are displayed. (B) CRISPR/Cas9 homozygous frameshift mutation in *Cbl* was created in RAW264.7 and tested with the *lpqN luxBCADE* strain as described for (A). The assay was performed three times with four independent mutant clones and two independent wild-type control clones. Each condition was measured in quadruplicate; mean ± SEM from representative clones are displayed. (C) *Cbl*^-/-^ and *Cbl*^+/+^ primary BMMs were infected with the *lpqN*, *luxBCADE* mutant and bacterial growth was enumerated by plating for CFU 12 days post-infection. (D) Growth of the *lpqN* mutant was also assessed in the *Cbl*^-/-^ and *Cbl*^+/+^ primary BMMs as described in (C) by measuring luminescence. Mean ± SEM are displayed. (E) The *Cbl* CRISPR/Cas9 clones of RAW264.7 described in (B) were infected with *L. monocytogenes* and CFU enumerated at the indicated times. Mean change in CFU relative to t=0 is shown ± SEM. Representative results from two independent experiments are shown. (F) Infection as in (E) with S. Typhimurium.

**Figure S4.**
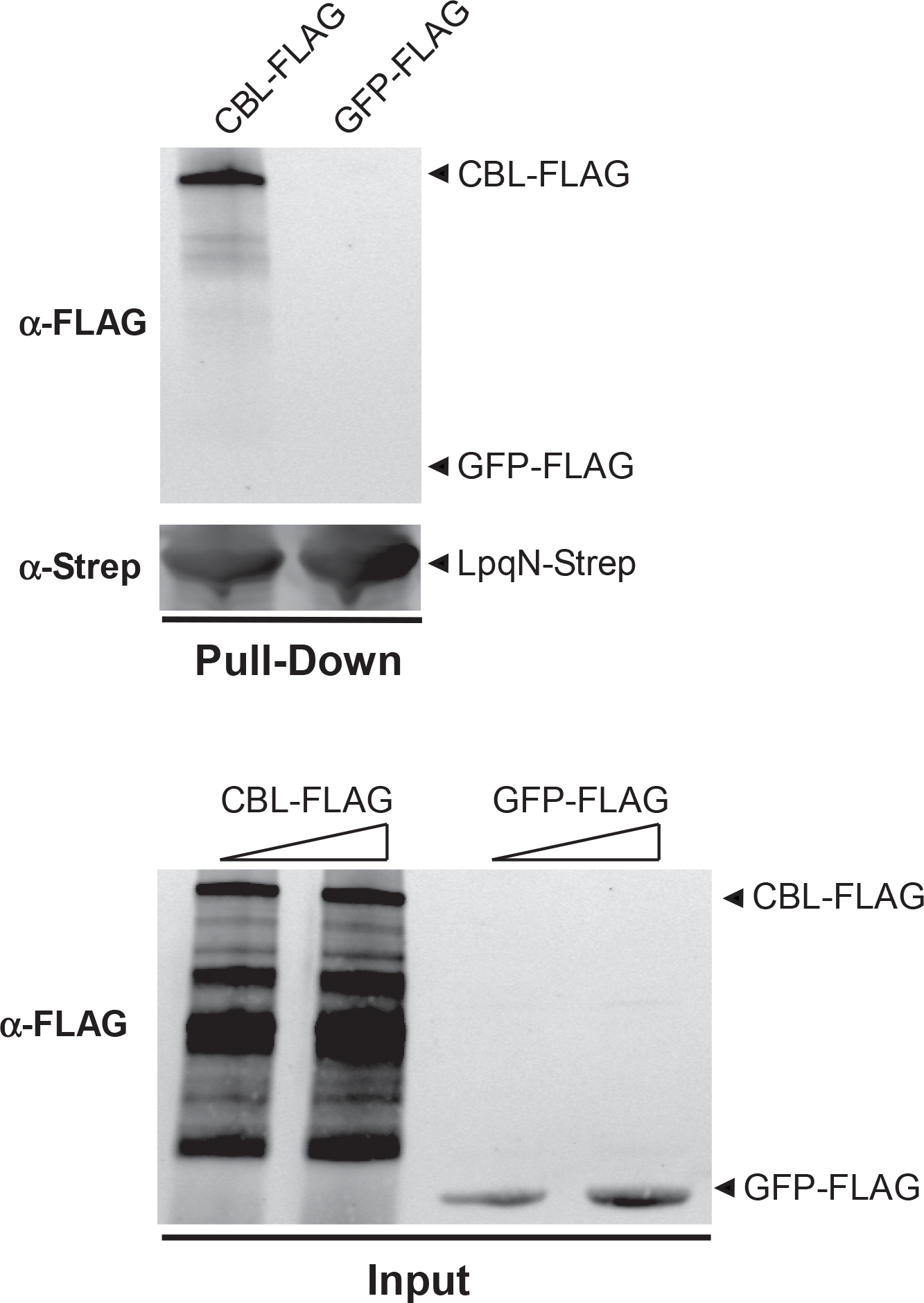
Related to Figure 4. Direct interaction between LpqN and CBL *in vitro*. LpqN-Strep was expressed in *E. coli* BL21DE3 cells, and whole cell lysates were mixed with similar lysates from cells expressing either CBL-FLAG or GFP-FLAG for 3h at 4°C to allow for binding. LpqN-Strep was affinity purified using Strep-Tactin resin, and co-purification of the host fusion proteins were detected by western blotting using anti-FLAG antibodies.

**Table S1. Related to Figure 1. High-confidence set of *Mtb* secreted proteins.**

*Mtb* culture filtrates separated by SDS-PAGE (Figure S1A) were subjected to LC-MS/MS analysis to identify secreted proteins. These results were manually curated to remove known cytoplasmic or cell-wall contaminants (Målen et al., 2007).

**Table S2. Related to Figure 1. Comparison of U937 and 293T co-purifying proteins.**

To evaluate the impact of addition of U937 macrophage cell lysates to our PPI scheme (Figure 1A), a subset of bacterial proteins were re-analyzed by AP-MS using 293T lysate alone without inclusion of the U937 lysate. All these interactions were scored by MiST and those with values ≥0.7 are displayed.

**Table S3. Related to Figure 1. Full AP-MS dataset.**

All host proteins identified from the AP-MS proteomics pipeline described in this study (Figure 1).

**Table S4. Related to Figure 1. MiST scoring of AP-MS dataset.**

**Table S5. Related to Figure 2. Comparison of AP-MS datasets from different pathogens.**

**Table S6. Related to Figure 2. Functional annotation of high-confidence interacting host proteins.**

DAVID v6.8 Uniprot keywords analysis was used to annotate all host proteins with MiST ≥0.7 for the indicated pathogens.

**Table S7. Related to Figure 2. Evolutionary analysis of *Mtb*-interacting proteins.**

SnIPRE and iHS analysis data.

**Table S8. Related to Figure 4. PTMs of host proteins during infection with *Mtb*.**

## STAR METHODS

### KEY RESOURCE TABLE

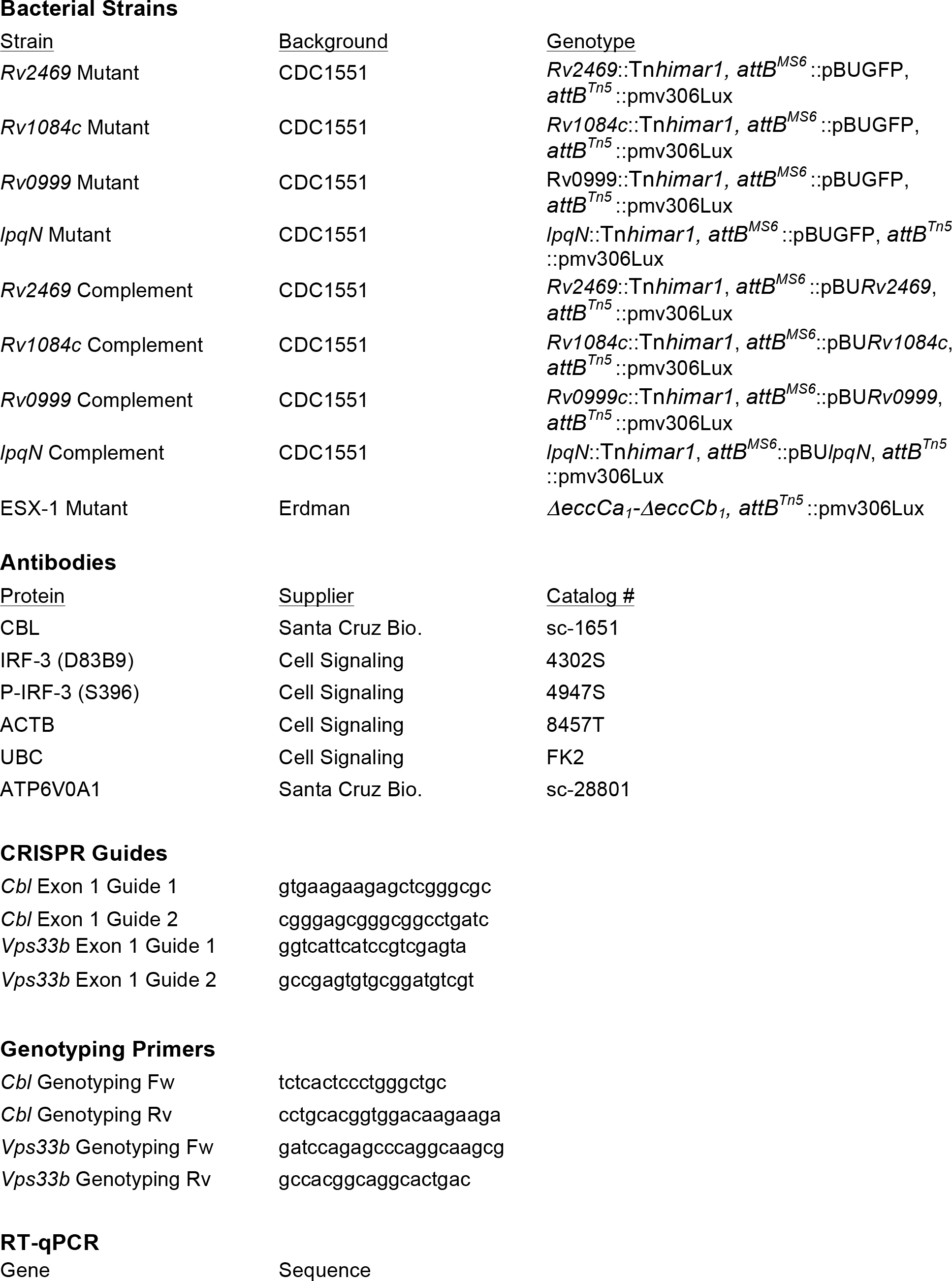

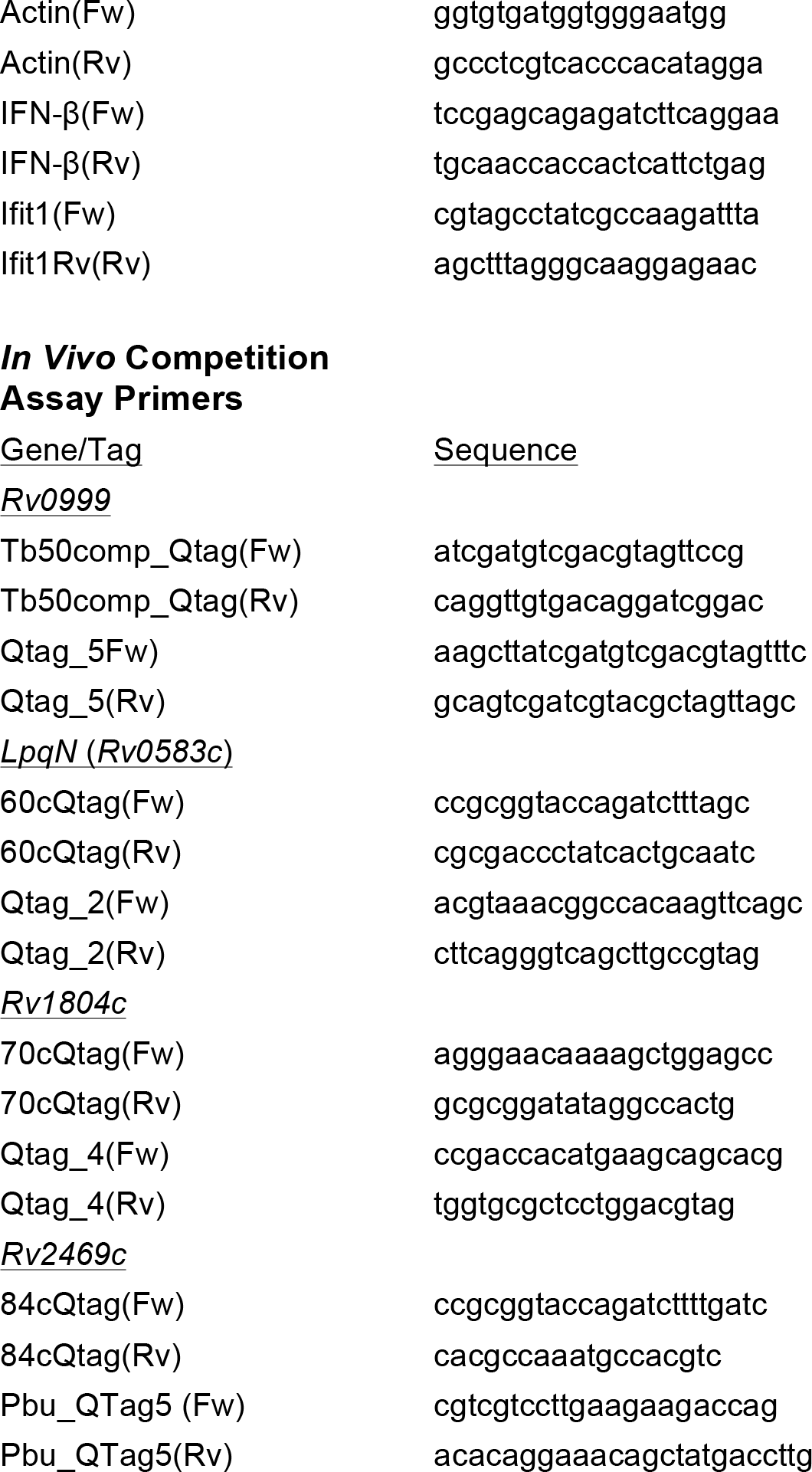

## CONTACT FOR REAGENT AND RESOURCE SHARING

Further information and requests for resources and reagents should be directed to and will be fulfilled by the Lead Contact, Jeffery Cox.

## EXPERIMENTAL MODEL AND SUBJECT DETAILS

### Macrophages

Bone marrow-derived macrophages (BMMs) were isolated by flushing the femurs from 8-10-week-old B6 female mice. They were cultured in high-glucose DMEM supplemented with 20% heat-inactivated FCS and 10% conditioned media from 3T3-MCSF cells. All cells were cultured with at 37°C with 5% CO_2_. *Cbl*^+/+^ and *Cbl*^-/-^ BMMs were isolated from mice of the following genotypes: 
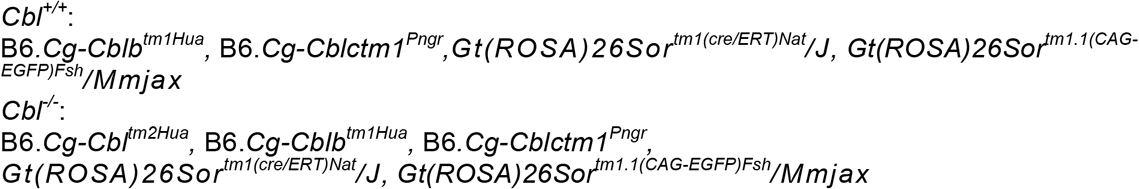

Both *Cbl*^+/+^ and *Cbl*^-/-^ BMMs were then cultured in the presence 1uM 4-hydroxytamoxifen (Sigma) to trigger recombination of the floxed *Cbl* allele (Mukhopadhyay et al., 2016).

### Cell Lines

RAW 264.7 mouse macrophages and 293T cells were purchased from ATCC and cultured according to ATCC recommendations with addition of 20mM HEPES pH7.4. 293T were purchased from ATCC and cultured according to ATCC recommendations.

### Bacterial strains

All *Mtb* strains were cultured in 7H9 liquid media (BD) supplemented with 10% Middlebrook OADC (Sigma), 0.5% glycerol, 0.05% Tween80 in roller bottles at 37°C. Transposon mutants were obtained from ATCC and carry *Himar1* transposon insertions with the Kanamycin resistance gene in the indicated locus. Transposon insertion sites were validated by PCR and Sanger sequencing (Lamichhane et al., 2003). For genetic complementation studies, the predicted promoter region and open reading frame were cloned into the SacI site of the integrating vector pBU (Vultos et al., 2006) which had been modified to confer Zeocin resistance, and strains were selected with 25ug/ml Zeocin. Control strains carried pBU with a ~100 nucleotide fragment of GFP as unique sequence tag. For luminescent growth assays, strains carried the containing a codon-optimized *luxBCADE* operon expressed from a MOPS promoter carried on the pmv306-Hyg integrating vector (Craney et al., 2007). The *ΔeccCa*_1_-*ΔeccCb*_1_ strain was made by specialized transduction using the pMSG361 vector (Rosenberg et al., 2015).

## METHOD DETAILS

### *Mtb* secreted protein analysis

*Mtb* Erdman strain was grown in 7H9+OADC (BD Biosciences), +0.05% Tween-80 (Sigma) until mid-log, then transferred to Sauton’s minimal media + 0.05% Tween-80 for 5d, and then finally to Sauton’s media with 0.005% Tween for 5d. Bacteria were pelleted and supernatants concentrated with a 3K MWCO filter (Millipore). Bacteria were lysed by boiling, followed by 10 minutes of sonication in 1% SDS. 20ug each of cell lysate and culture filtrate were separated by SDS-PAGE and western blot performed against GroEL to verify that there was no detectable contamination of the culture filtrate by cytoplasmic proteins. For MS analysis, protein was resolved on SDS-PAGE, visualized by Coomassie, and gel slices subjected to trypsin digestion as described below.

### *In vivo* competition assay

12-week-old female B6 mice were inoculated by aerosol with a pool of four mutant strains and cognate complemented strains delivered at ~75 CFU of each strain using a Madison chamber device. At the indicated times, lungs of 4-5 mice were homogenized and plated on 7H10 plates. ~10,000 individual colonies were scraped into Trizol (Thermo Fisher Scientific). Samples were then lysed using silica beads and bead-beating and RNA removed from Trizol per manufacturer instructions for RNA isolation. Genomic DNA was then isolated from organic phase by back-extraction of with 4 M guanidine thiocyanate/50mM Sodium Citrate/1M Tris base, and the aqueous phase containing DNA was removed. The pH was normalized by adding 0.2 volumes sodium acetate, and 0.4 volumes ethanol was added. DNA was then purified over RNeasy columns (Quiagen) according to manufacturer instructions. qPCR primers were designed to detect unique sequence tags inserted in each strain’s genome using either a fragment of the GFP gene (mutant strains) or the unique junction between the pBU vector & complementation cassette (complemented strains). Each primer set was verified to detect only its specific strain. qPCR was conducted with Taq polymerase (NEB) and SYBR green I (Sigma) detection.

### *Ex vivo* luminescent bacterial growth assay

*Mtb* strains carrying the *luxBCADE* operon were prepared for inoculation by washing twice in PBS, removing aggregates with a 200 RCF spin, followed by gentle sonication to generate a fine bacterial suspension. Bacteria were opsonized in 10% heat-inactivated horse serum and macrophages infected at an MOI=2 by spinning inoculated plates for 10 min at 400 RCF. Monolayers were washed, and bacterial luminescence measured at the indicated times using a Spectromax L (Molecular Dynamics). Macrophage growth media was changed daily for cell lines and every 48h for BMM cultures.

### CRISPR mutagenesis

CRISPR guides were designed to target regions in the first exon of target genes using an online bioinformatic tool (http://crispr.mit.edu), and cloned into the BsmB1 site of the pXPR_001 vector (a gift from Feng Zhang (Addgene plasmid # 52961)). Viral particles were produced in Lenti-X cells (Clontech) per manufacturer’s instructions, and used to transduce RAW264.7 cells. Therese were selected in 5ug/ml Puromycin (InVivoGen), and single-cell clones isolated. Mutations were identified by using PCR to amplify the first exon by PCR followed by Sanger sequencing. CRISPR CBL mutants were independently confirmed by western blot with a mouse anti-c-CBL antibody (Santa Cruz Biotechnology cat# sc-1651).

### Western blots

Protein in lysates was quantified by BCA (Pierce Fisher Scientific). 20ug of cell lysate was separated by SDS-PAGE (Bio-Rad TGX), and transferred onto nitrocellulose membranes. After probing with the indicated antibodies, membranes were then imaged on an Odyssey scanner (Li-cor).

### Affinity purification

*Mtb* genes were amplified by PCR from Erdman strain genomic DNA. For genes with a predicted signal peptide using SignalP 3.0 (Bendtsen et al., 2004), the portion of the open reading frame corresponding to the mature protein was amplified. Genes were cloned into PCDNA4 with a C-terminal 2×-Strep tag and expressed in 293T cells using calcium phosphate transfection. Cells were lysed 36h later in IP buffer (50mM Tris 7.4, 150 mM NaCl, 1mM EDTA, 0.05% NP-40) with phosphatase and protease inhibitors (Roche) and tagged proteins immobilized on Strep-Tactin resin (IBA). Macrophage lysate was generated from U937 cells differentiated with 10nM PMA for 72h and then similarly lysed in IP buffer. 10mg of macrophage lysate was added to each immobilized bacterial factor and binding allowed to proceed overnight. The resin was then washed 4× in IP buffer, and twice in IP buffer without NP-40 before elution in 10mM biotin. For reciprocal immunoprecipitation using FLAG-tagged human proteins the human factors were cloned as 3×FLAG-tag fusion proteins in pCDNA4. Combinations of host and bacterial factors were co-transfected into 293T cells using calcium phosphate and immunoprecipitation with anti-FLAG M2 antibody was performed using the same lysis and wash conditions as above.

### Mass Spectrometry

Purified proteins eluates were digested with trypsin for LC-MS/MS analysis. Samples were denatured and reduced in 2M urea, 10 mM NH4HCO3, 2 mM DTT for 30 min at 60C, then alkylated with 2 mM iodoacetamide for 45 min at room temperature. Trypsin (Promega) was added at a 1:100 enzyme:substrate ratio and digested overnight at 37C. Following digestion, samples were concentrated using C18 ZipTips (Millipore) according to the manufacturer’s specifications. Desalted samples were evaporated to dryness and resuspended in 0.1% formic acid for mass spectrometry analysis.

For the MS study to establish a high-confidence set of *Mtb* secreted proteins culture filtrates were resolved by SDS-PAGE. The gel was partitioned into sequential slices and each of these was individually subjected to in-gel trypsin digestion followed by desalting and LC-MS/MS analysis on a Thermo Scientific LTQ XL linear ion trap mass spectrometer. The LTQ XL system was equipped with a LC Packings Ultimate HPLC with an analytical column (10 cm × 75 um I.D. packed with ReproSil Pur C18 AQ 5 um particles). The system delivered a gradient from 5% to 30% ACN in 0.1% formic acid over one hour, and collected data in a data-dependent fashion. The LTQ XL collected one full scan followed by 10 collision-induced dissociation MS/MS scans of the 10 most intense peaks from the full scan. Dynamic exclusion was enabled for 30 seconds with a repeat count of 1.

For the primary AP-MS study used to establish the interactome, digested peptide mixtures were analyzed by LC-MS/MS on a Thermo Scientific LTQ XL linear ion trap mass spectrometer as above.

For the secondary analysis, where AP-MS was performed without inclusion of U937 macrophage lysate, the digested peptide mixtures were analyzed by LC-MS/MS on a Thermo Scientific Velos Pro dual linear ion trap 238 mass spectrometer equipped with an Easy-nLC II HPLC with a pre-column (2 cm × 100 um I.D. packed with 239 ReproSil Pur C18 AQ 5 um particles) and an analytical column (10 cm × 75 um I.D. packed with ReproSil Pur 240C18 AQ 3 um particles). A gradient was delivered from 5% to 30% ACN in 0.1% formic acid over one hour. 241. The mass spectrometer collected data in a data-dependent fashion with one full scan followed by 20 collision-242 induced dissociation MS/MS scans of the 20 most intense peaks from the full scan. Dynamic exclusion was 243 enabled for 30 seconds with a repeat count of 1.

The results raw data was matched to protein sequences by the Protein Prospector algorithm (Clauser et al., 1999). Data were searched against a database containing Swiss Prot Human protein sequences (downloaded March 6, 2012), and concatenated to a decoy database where each sequence was randomized in order to estimate the false positive rate. The searches considered a precursor mass tolerance of 1 Da and fragment ion tolerances of 0.8 Da, and considered variable modifications for protein N-terminal acetylation, protein N-terminal acetylation and oxidation, glutamine to pyroglutamate conversion for peptide N-terminal glutamine residues, protein N-terminal methionine loss, protein N-terminal acetylation and methionine loss, and methionine oxidation, and constant modification for carbamidomethyl cysteine. Prospector data was filtered using a maximum protein expectation value of 0.01 and a maximum peptide expectation value of 0.05.

Our global identification of phosphorylation and ubiquitylation during *Mtb* infection of RAW264.7 macrophages is described in a separate manuscript (Parry et. al., *in preparation*) and will be submitted to ProteomeXchange via PRIDE.

### MiST

AP-MS samples were scored with Mass spectrometry Interaction STatistics (MiST) algorithm, using the MiST reproducibility (0.45), specificity (0.50) and abundance (0.05) weights previously published (Davis et al., 2015). All bait-prey pairs with a MIST score ≥ 0.70 were considered confident interactions and were combined with human protein interactions from the CORUM and STRING databases. The resulting network diagram was plotted using Cytoscape, v.3.1.2 411 (Smoot et al., 2011). For the subset re-analysis without U937 lysate, MiST with identical settings was used but additional control Strep-tag purifications added to the analysis matrix in MiST to compensate for the smaller number of experimental samples and establish robust specificity scoring.

### Evolutionary Analysis

For analysis over a 5-7 million-year timescale we employed SnIPRE (Eilertson et al., 2012) which is a Bayesian mixed effects model of the McDonald-Kreitman test (McDonald and Kreitman, 1991). We focus our analysis on the human lineage by removing sites that are inferred to have received mutations along the chimpanzee lineage based on an alignment of closely related primates. For analysis of more recent evolution in the primate lineage we used a modified version of integrated Haplotype Score (iHS) (Voight et al., 2006) as implemented in (Szpiech and Hernandez, 2014).

### In *Vitro* Interaction Studies

Proteins were cloned into the pH3C vector as either a Strep-Tag fusion (LpqN) or FLAG-Tag fusions (CBL, GFP). Factors were individually expressed in BL21 cells by overnight induction with IPTG at room-temperature. Cells were lysed in PBS + protease inhibitors using lysozyme and sonication. Total cell lysates containing ~10ug of each tagged protein were combined, subjected to Strep-Tactin purification, and interacting proteins were detected by western blotting as above.

### Gene Expression Analysis

Infected macrophages were lysed in Trizol (Thermo Fisher Scientific) and then purified with silica spin-columns (Purelink, Ambion) per manufacturer instructions. cDNA was generated using 500ng total RNA with the Superscript III First Strand Synthesis Kit (Invitrogen), and subjected to qPCR as described above with normalization to the beta-actin transcript.

### Immunofluorescence microscopy

Coverslips were fixed for 20min in 4% PFA, permeabilized in 0.05% Saponin (Sigma) and incubated with the indicated antibodies for 3h at RT. Mycobacteria were visualized either by expression of mCherry or by labeling with SytoBC (Fisher). All samples were mounted and images acquired on a Keyence BZ-X700 microscope. Z-stacks at 0.5uM were acquired and maximum intensity projections generated. To quantify colocalization positive phagosomes were scored for each marker by a microscopist blinded to sample identity.

## QUANTIFICATION AND STATISTICAL ANALYSIS

### Statistical Analysis

For RT-qPCR, each sample was amplified in triplicate and transcript levels normalized to the beta-actin gene and the standard error of the mean was calculated. Each experiment was repeated at least two times from separate biological samples with representative data from one biological replicate shown. For luminescent growth assays each sample was quantified in quadruplicate wells and standard-error of the mean was calculated. Each experiment was repeated at least three times from separate biological samples with representative data from one biological replicate shown. For *in vivo* competition assays the sequence tags were amplified in triplicate qPCR reactions for each mouse, and the ratio between mutant and complement calculated. For the set of mice at each time point, the mean mutant:complement ratio and standard-error of mean were calculated. A Kruskal-Wallis test was used to calculate p-values for comparisons among sets of mice.

### Bioinformatic Analysis of Host Proteins

The set of high-confidence host interacting proteins was analyzed using the DAVID Bioinformatics Resources 6.8 functional annotation tool. Uniprot accession numbers were matched with Uniprot keyword categorization to map protein function and determine enrichment relative to the human proteome. Thresholds for inclusion were set to two-fold enrichment with a Benjamini–Hochberg corrected p-value of <0.05 to adjust for multiple hypotheses testing. Categories were manually curated to remove redundant and nonspecific categorizations. For overlap analysis between datasets a Fisher’s exact test was used with the background proteome defined empirically as the set of all proteins detectable by our Velos Pro dual linear ion trap instrument in 293T cell-related experiments (10,782 proteins)

## DATA AND SOFTWARE AVAILABILITY

The MiST algorithm is available freely on github (https://github.com/kroganlab/mist), and the Comppass algorithm at (https://github.com/RGLab/COMPASS). All MS data will be deposited in the PRIDE database

